# A mesocosm experiment in ecological physiology: adaptive modulation of energy budget in a hibernating marsupial under chronic caloric restriction

**DOI:** 10.1101/2020.06.05.136028

**Authors:** Roberto F. Nespolo, Francisco E. Fontúrbel, Carlos Mejias, Rodrigo Contreras, Paulina Gutierrez, José Ruiz, Esteban Oda, Pablo Sabat, Catherine Hambly, John R. Speakman, Francisco Bozinovic

**Affiliations:** Center of Applied Ecology and Sustainability (CAPES), Departamento de Ecología Facultad de Ciencias Biológicas, Pontificia Universidad Católica de Chile, Santiago, Chile; Instituto de Ciencias Ambientales y Evolutivas, Universidad Austral de Chile, Valdivia, Chile; Millennium Institute for Integrative Biology (iBio), Santiago, Chile; Instituto de Biología, Pontificia Universidad Católica de Valparaíso, Valparaíso, Chile; Departamento de Ciencias Ecológicas, Facultad de Ciencias, Universidad de Chile, Santiago, Chile; Institute of Biological and Environmental Sciences, University of Aberdeen, Aberdeen, AB24 2TZ, UK; State Key Laboratory of Molecular Developmental Biology, Institute of Genetics and Developmental Biology, Chinese Academy of Sciences, Beijing, 100101, China; Chinese Academy of Sciences Center of Excellence in Animal Evolution and Genetics, Kunming, China

**Keywords:** behavioral thermoregulation, chronic caloric restriction, daily energy expenditure, doubly labelled water, energy budget, hibernation, marsupial, social clustering

## Abstract

During the last sixty years, mammalian hibernation (i.e., seasonal torpor) has been interpreted as a physiological adaptation for energy economy. However -and crucially for validating this idea – direct field comparisons of energy expenditure in hibernating and active free-ranging animals are scarce. Using replicated mesocosms and a combination of energy budgeting approaches (i.e., doubly labelled water, rates of CO_2_ production and food intake), we experimentally manipulated energy availability and quantified net energy costs of hibernation in a marsupial. We hypothesized that, when facing chronic caloric restriction (CCR), a hibernator should maximize torpor use for compensating the energetic deficit, compared to *ad libitum* fed individuals (=controls). However, intensifying torpor duration at low temperatures could increase other burdens (e.g., cost of rewarming, freezing risk). In order to explore this trade-off, we followed the complete hibernation cycle of the relict marsupial *Dromiciops gliroides*, and estimated its total energy requirements, and compared this with a control condition. Our results revealed: (1) that energy restricted animals, instead of promoting heat conservation strategies during hibernation (e.g., social clustering and thermoregulation), maximized torpor use and saved just enough energy to cover the deficit, and (2) that hibernation represents a net energy saving of 51% compared with animals that remained active. This work provides compelling evidence of a fine-tuning use of hibernation in response to food availability and presents the first direct estimation of energy savings by hibernation encompassing the total hibernation cycle.

## Introduction

The countless ways natural selection shapes organismal design and function has always intrigued biologists, particularly in ecosystems where energy availability is diluted, temporally or spatially variable (Mueller and Diamond 2001, Ferguson 2002, Nie et al. 2015). In this scenario, energy flow is often explained by the allocation principle, where energy from food passes through several sequential bottlenecks (e.g., foraging, digestion, assimilation), and must be allocated to different functions in parallel (e.g., growth, maintenance, and reproduction) (Weiner 1992). From this perspective, nature’s economy would be defined by austerity, for which ectotherms provide the best fit to the rule, as they minimize maintenance costs when activity is low (Pough 1980, Artacho and Nespolo 2009). Endotherms (birds and mammals) on the opposite have a wasteful lifestyle, a counter-intuitive solution for any idea of nature’s economy (an “extravagant economy” *sensu* (Hayes and Garland 1995, Koteja 2004). However, some endotherms experience transient periods of ectothermy or torpor (=heterothermy, hereafter), as putative adaptations to seasonal or unpredictable reductions in environmental productivity. For the case of hibernation (i.e., seasonal multi-day episodes of torpor)(Geiser and Ruf 1995b), animals experience drops in body temperature and a general reduction in metabolism lasting several days or weeks, where body temperature is maintained a few degrees above ambient temperature. During these episodes, maintenance costs fall to a fraction of normal values, with significant energy savings (Geiser 2004), a “logical” solution for animals that cannot migrate to better environments (Schmidt-Nielsen 1979). Thus, hibernators would have the long-term benefits of endothermy, together with the short-term benefits of ectothermy.

Contrarily with daily torpor, where metabolic depression occurs during a few hours, hibernation is characterized by torpor events that increase in duration and frequency as the cold season progresses {Geiser, 2013 #10429}. Thus, animals modulate the frequency of such events depending on the cold, photoperiod and the amount of fat reserves, the latter being determinant on predicting hibernation survival (Humphries et al. 2002, Humphries et al. 2003a, Humphries et al. 2003b). But how much energy, exactly, is saved during a complete hibernation cycle, compared to a situation without hibernation? Do hibernating animals regulate torpor frequency “wisely” as food availability varies? Although hundreds of laboratory experiments have provided partial answers to these questions, only a handful of experimental manipulations of food availability have demonstrated a link between energy availability, torpor frequency and fat reserves in hibernation (reviewed in (Vuarin and Henry 2014).

According to Boyles et al. (Boyles et al. 2020), to compensate for reduced energy availability, a hibernator that perceives an energetic bottleneck in the environment should experience longer and deeper torpor bouts and select sites with low temperatures for hibernating (Song et al. 2000). However, this has a limit imposed by several costs (e.g., prolonged inactivity, freezing mortality, decreasing immune function and sleep deprivation, see Humphries et al. 2003b, Boyles et al. 2020), which furnishes a “hibernation trade-off” where an optimum (minimum) hibernation temperature is defined {Humphries, 2002 #10368}. Above this temperature, energy saved by hibernation is maximized and below this temperature, hibernation costs are maximized. In nature, a range of responses have been observed. For instance, passerine birds (Wojciechowski et al. 2011, Douglas et al. 2017), mice (Eto et al. 2015) and Siberian hamsters (Jefimow et al. 2011) minimize heat loss during daily torpor, whereas non-migrating bats (Ryan et al. 2019) and sugar gliders (Nowack and Geiser 2016) minimize body temperature during multi-day hibernation.

Here we explored the hibernation trade-off on the social Microbiotheriid marsupial *Dromiciops gliroides* (Hershkovitz 1999) using a mesocosm setup for tracking animals during a complete hibernation cycle. Specifically, we manipulated food by applying a chronic caloric restriction treatment (CCR) and we measured total energy requirements for wintering using gross energy intake (=daily food consumption) and CO_2_ production, using the doubly labelled method. Specifically, we predicted that CCR animals (compared with ad libitum fed animals) will either intensify torpor use in order to maximize energy savings and compensate for the energy restriction they will avoid risks by using heat conservation strategies (e.g., social clustering and hibernacula use).

## Methods

### Animals

*Dromiciops gliroides* (Thomas 1894) is the only living species of Microbiotheria; the ancestral group of Australian marsupials. *D. gliroides* is a small arboreal marsupial inhabiting the temperate rainforests of southern South America, living in native forest stands dominated by *Nothofagus* spp. and *Araucaria araucana* trees (Hershkovitz 1999, Fonturbel et al. 2012). This marsupial is known to be the sole disperser of several endemic plant species, thus being intimately associated with the temperate rainforest (Amico et al. 2009), where this experiment was performed. We installed the mesocosm in Estación Experimental Fundo San Martin (SM), a property of Universidad Austral de Chile (39° 41’S 73° 18’W), whichh is within the typical habitat of *Dromiciops gliroides*. In this paper we refer to “hibernation” as the multiday torpor bouts lasting several days, in contrast to daily heterotherms that a experiences torpor bouts of 3-12 hours (Geiser and Ruf 1995a). No previous monitoring of the whole hibernation period of *D. gliroides* is available, which was estimated to extend from May to September (Hershkovitz 1999, Muñoz-Pedreros et al. 2005). Thus, we started the experiment in April, and finished data gathering in December. We captured 40 individuals from different sites within SM during the austral summer, which were were live-captured using Tomahawk-like traps baited with banana and attached to the trees, 2 m above the ground (Fonturbel 2010). Traps were located 300 m apart from the enclosure site, in four different patches of forest, each on a sampling grid. Each individual was marked using PIT-tag (BTS-ID, Sweden) subcutaneous mark, and transported to the laboratory inmmediately after capture for feeding and rehydration.

### Outdoor enclosures

To characterize simultaneosly physiological and thermoregulatory responses of hibernating *D. gliroides*, we built eight cylindric enclosures (Fig. 1), which were distributed within the forest and separated about 5 m from each other, covering a total area of about 80 m^2^ (see Supplementary Material). Each enclosure had a internal volume of 2 m^3^, and was manufactured in zinc with a large 1.8m-diameter cylinder buried 10 cm in the ground, which gave a 0.8 m height above ground. Each enclosure had a data logger installed for continuous measurement of air temperature (HOBO ®). Initially, four enclosures were assigned to a control treatment (”control”, hereafter) and the other four were assigned to a caloric restriction treatment (”CCR”, hereafter; see below). Five unrelated animals (i.e., from different sites to avoid kinship effects) (Franco et al. 2011) were released in each enclosure, on April 1^st^ (autumn). Unfortunatelly, one of the CCR enclosures was destroyed by a tree falling during winter (animals escaped), which left us with an unbalanced design with 35 animals across 7 enclosures (4 controls and 3 CCR).

**Fig 1.**
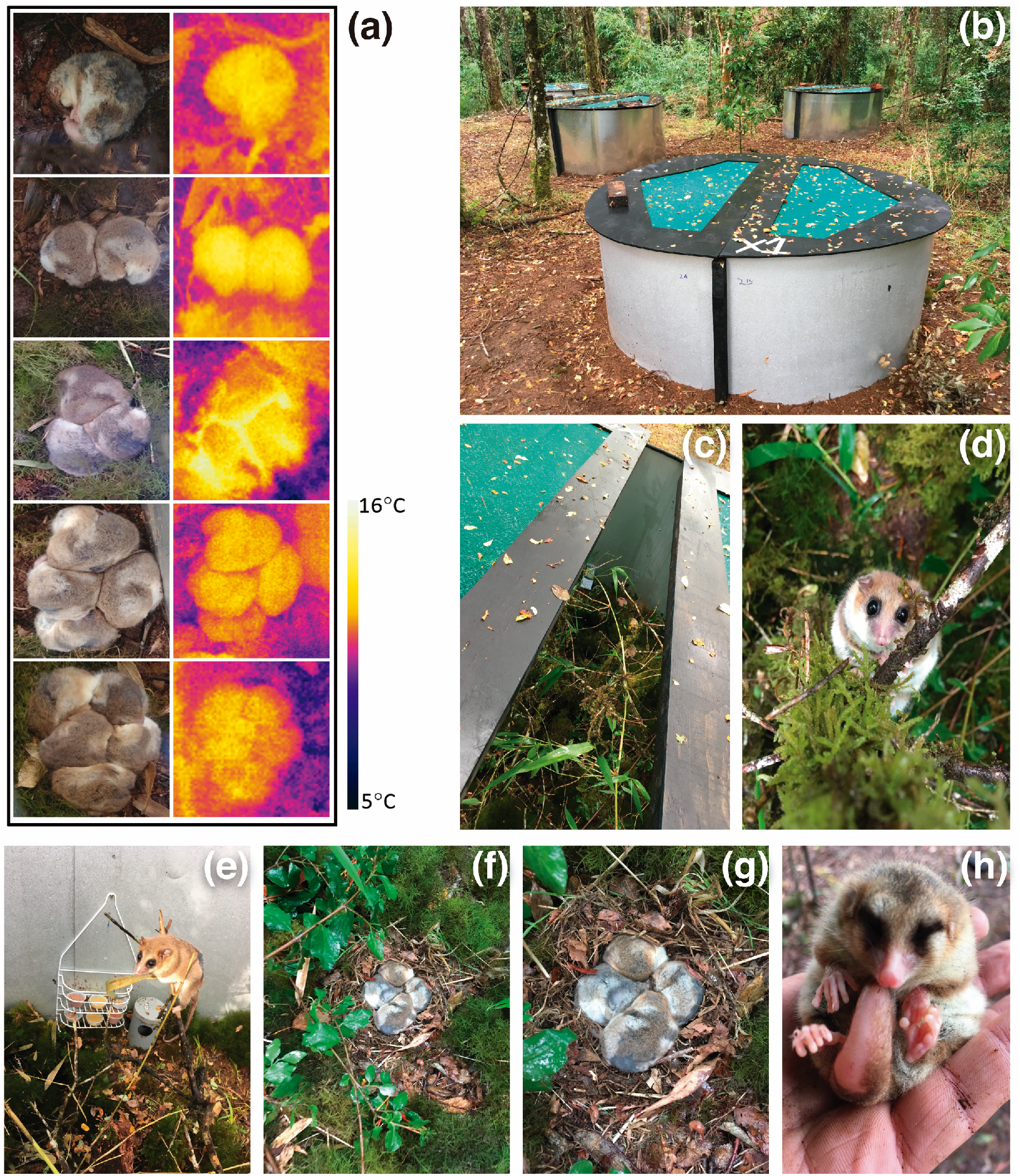
a) Digital photographs and thermographs of clustered hibernating *D. gliroides*, at different cluster sizes. The average temperature of each picture is 10°C, approximately. b) Photographs of the enclosures (c), enclosure opening showing the reproduced forest environment, (d) female *Dromiciops* within the enclosure, (e) a male with the food feeders, (f) a cluster of hibernating animals after removing the hibernaculum, (g) a close-up of a cluster of 5 hibernating animals (h) a torpid female of the control treatment. Red arrows indicate the moment of daily energy expenditure and basal metabolic rate measurements.

### Experimental energy manipulation

To explore how constant food shortage induce compensatory responses during hibernation, we applied a chronic caloric restriction treatment to three enclosures. Then, we offered the equivalent of 165 kJ ind^−1^d^−1^ for the control enclosures and provided to the CCR animals, 60% of this value (95 kJ ind^−1^d^−1^). The food was provided in equal volumes every day, but once a week we provided a fresh weighed amount (± 0.01 g) to each enclosure and weighed the fresh weight of the leftovers for drying to constant weight (60°C). With this, we estimated the water content of the diets for estimating average energy intake. Using weekly values of energy consumption, we calculated the (per capita) total hibernation energy requirements (kJ per individual).

### Torpor thermoregulation and daily energy expenditure

Weekly, we took digital thermographic images to clustered torpid individuals in order to estimate the thermal differential between animals and substrate and to relate this to the caloric restriction treatment (Fig 1a). We also recorded cluster sizes and whether animals were within or outside the hibernaculum (see Supplementary Material). To determine direct daily energy expenditure (DEE, kJ/day), we applied the doubly labelled water technique (Lifson and McClintock 1966, Butler et al. 2004)(see Supplementary Material) on 24 captive individuals before the release on enclosures (week zero, in summer, indicated in Fig 2a), we successfully repeated these determinations in 16 animals at week 18 of the experiment (late winter; eight individuals from the CCR treatment and eight from the control treatment), thus giving an average DEE for 48 hours. Basal metabolic rate (BMR) was determined from the rate of CO_2_ production in these same animals measured in the laboratory using standard respirometry techniques (Nespolo et al. 2010, Contreras et al. 2014).

**Fig 2.**
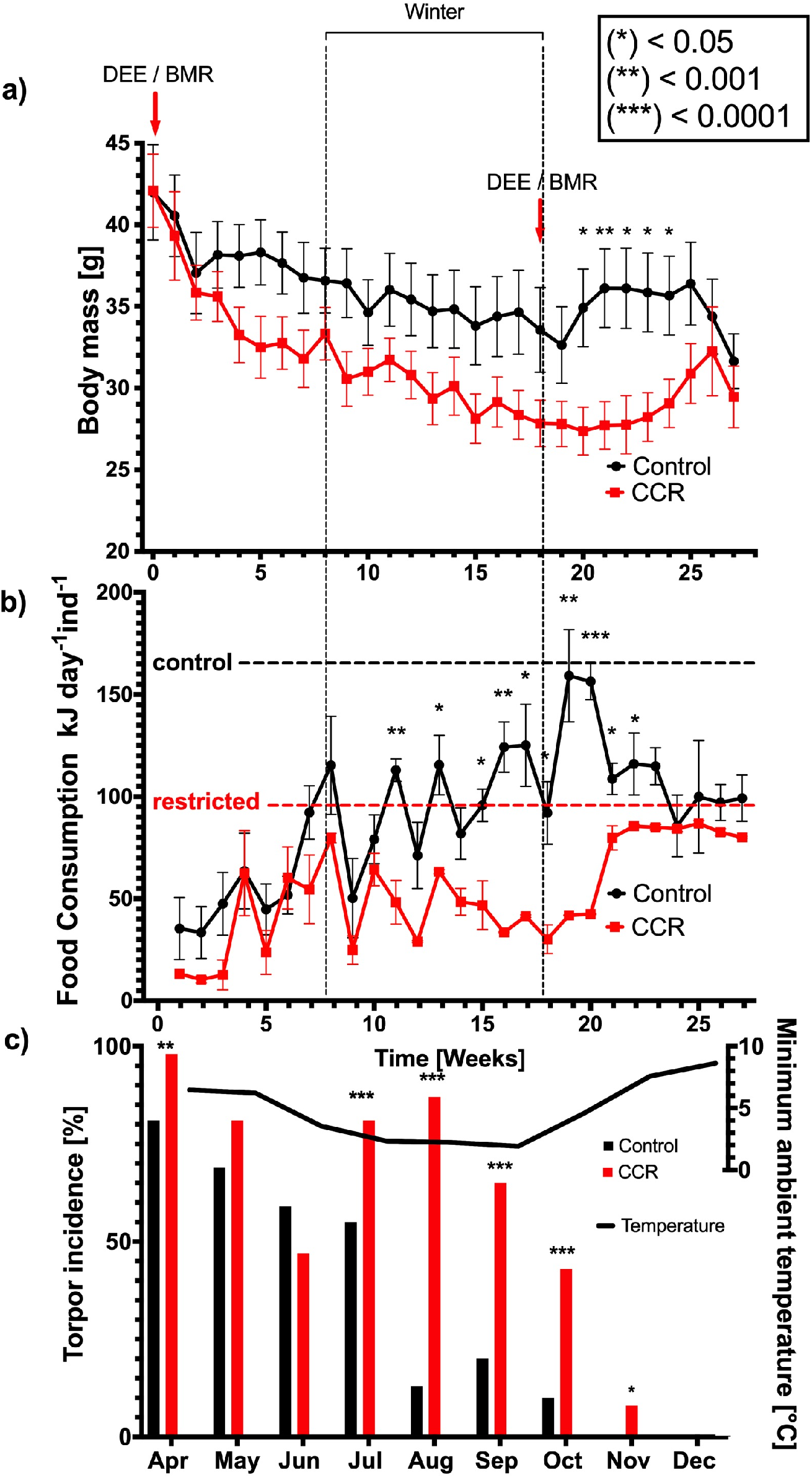
a) Weekly body masses (mean±sem) of individuals of *D. gliroides* either receiving food ad libitum or exposed daily to a chronic energetic restriction, CCR, since week 0 (April, 15^th^, autumn), in a semi-natural experiment (enclosures). Comparisons between CCR (n=15) and control (n=22) individuals were significant between week 20 and week 25 (t-tests, p<0.05); b) Per-capita energy consumption (dry mass) showing control (offered: 165 kJ ind^− 1^ day^−1^) and CCR (offered: 95 =kJ ind^−1^ day^−1^; indicated by horizontal dotted lines); c) Torpor incidence in CCR and control individuals (bars) and weekly minimum ambient temperature (line).

### Statistical analysis

We used a combination of generalized linear mixed models and standard parametric analyses such as ANCOVA, ANOVA and linear regressions when justified. Detailed descriptions of statistical analyses are provided in Supplementary Material.

All procedures presented in this study were approved by the Chilean Agriculture and Livestock Bureau (SAG) permits No 4371/2019 and 3393/2019, and by the Bioethics Committee of the Austral University of Chile, resolution 313/2018 annex 2019.

## Results

The main outcome of this experiment supports the idea that hibernating *D. gliroides* modulated torpor use for saving energy and cover the energetic deficit imposed by caloric restriction (results summarized in Fig. 2 and Supplementary Table 2). Indeed, animals under CCR (n= 15) consumed similar amounts of food as controls initially, but approximately at the eleventh week they consumed significantly less food than controls (n=20)(Fig 2b). CCR animals did not prefer to cluster in larger groups or use hibernacula for heat maintenance, and no statistical differences in any thermoregulatory aspect of the comparison of CCR and controls groups were observed (see Figs S2 and S3 in Supplementary Material). Moreover, those individuals experienced a constant reduction of body mass (M_B_); to become significant at the 20^th^ week (Fig. 2a). At week 23, however, CCR animals started to recover M_B_ and were not significantly different from controls by week 25 (two-tailed t-tests, p<0.001; Fig. 2a), thus suggesting that they, without access to extra food, managed their energy budget more efficiently. This is confirmed by measurements of per-capita energy consumption, which shows CCR animals consistently ingested less food than controls, until the rise in ambient temperatures during the austral spring (Fig. 2b-c). Then, energy intake became significantly higher in control individuals compared to CCR individuals at week 10 until week 24 (two-tailed t-tests, p < 0.01; Fig 2b). This is explained by a higher incidence in torpor use in CCR animals compared to controls, a difference that was the largest during August, which suggests that the main trigger of torpor was body condition rather than immediate food availability (Fig. 2c). Control animals attained a maximum weight loss of 13.2 ± 5.1% (mean ± sem) by week 19, whereas CCR animals reached a weigh loss of 34.8 ± 3.1% by week 20. Also, daily energy intake was significantly correlated with air temperature in CCR animals (p<0.01, n=432) whereas this correlation was non-significant for control animals (p=0.08, n=652; Fig 3). Thus, although CCR animals had access to 95 kJ ind^−1^per day, they reduced energy consumption to about half of this value (=47.7 ± 3.9 kJ day^−1^ ind^−1^, week 8-18, n= 3 enclosures), which was significantly lower than that in controls (96.7 ± 7.3 kJ day^−1^ ind^−1^, week 8-18, n= 4 enclosures)(p ≪ 0.001, t-test). This allowed them to reduce total winter energy requirements (i.e., per capita, *E_w_*) to 46% of the controls (control: *E_w_* = 10,066 ± 593.9 kJ ind-1, n= 4 enclosures; CCR: *E_w_* = 4,583.8 ± 113.6 kJ ind^−1^, n=3 enclosures; p < 0.001, t-test).

**Fig 3.**
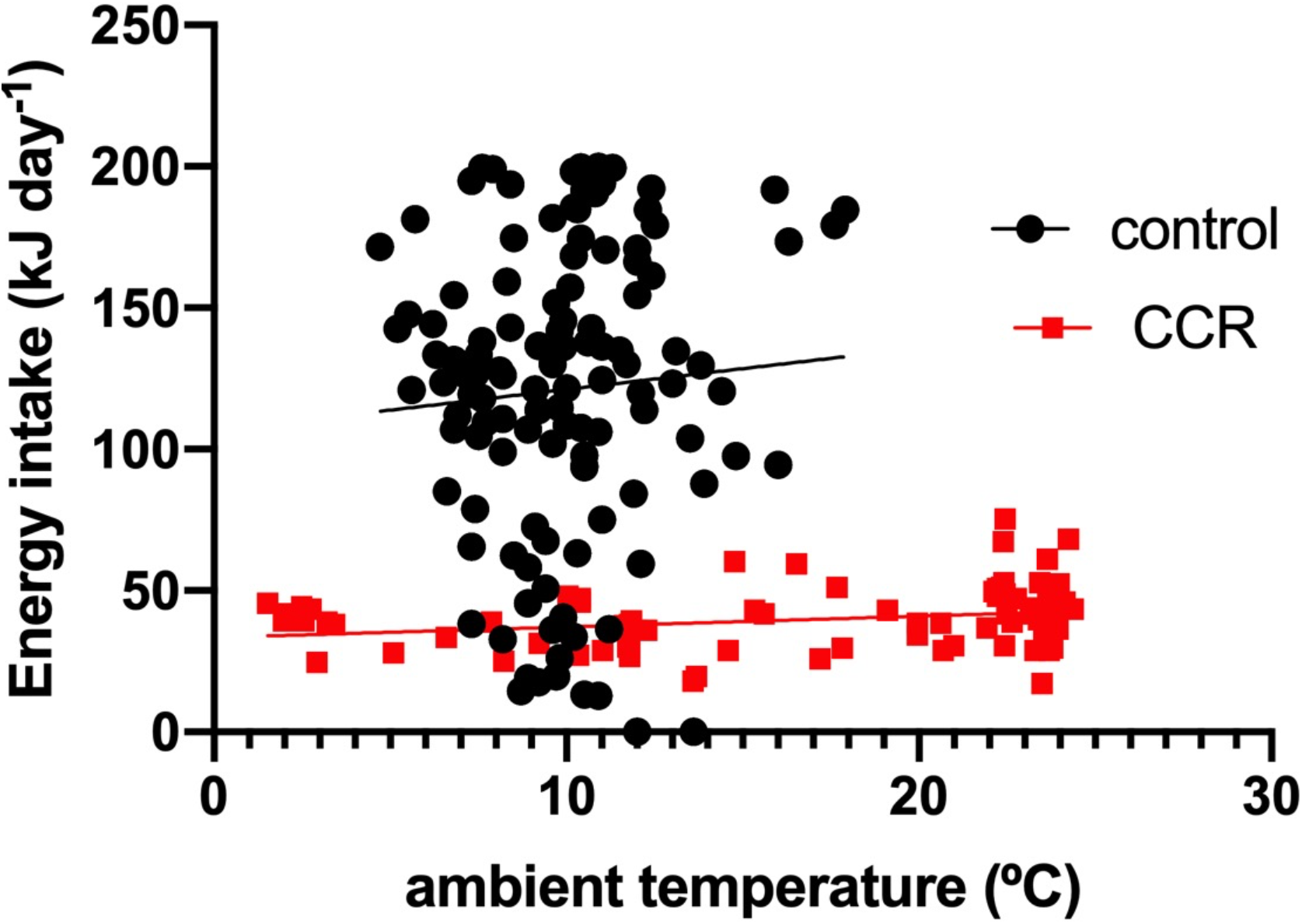
Daily energy intake estimated from food consumption in function of air temperature during the experimental period.

During the winter period (i.e., between weeks 8 to 18), animals exhibited an approximately constant negative slope in body mass (see Fig 2a). On average, each animal lost 3.0g (control) and 5.5g (CCR) in 70 days (i.e., 0.042 and 0.079 g day^−1^ind^−1^, respectively), which can be assumed to be 60% body fat (Mitchell et al. 2015). Thus, with an energy content of 39.7 kJ g^−1^ for fat (Walsberg and Wolf 1995), this gives 1.0 and 1.9 kJ day^−1^ ind^−1^, for each condition respectively. Thus, daily energy expenditure from food and body fat consumption can then be calculated as DEE = *E_w_* + *E_FAT_* in each case (being *DEE_CONTROL_* = *E_w-control_* + *E_FATcontrol_* and *DEE_CCR_* = *E_w-ccr_* + *E_FATccr_*). This gives: *DEE_CONTROL_* =98.6 kJ day^−1^ind^−1^ and *DEE_CCR_* = 48.7 kJ day^−1^ind^−1^. Thus, control animals, which were active at the moment of sampling, spent on average twice the amount of energy of CCR animals, which were in deep torpor.

The doubly labelled water measurements show that summer animals had a DEE of 44.9±2.2 kJ day^−1^ ind^−1^ (n=24) which is 58% of the expected DEE for mammals (Nagy 2005). This increased significantly in winter to 47.3 ± 5.6 kJ day^−1^ ind^−1^ (n=8) (82% of the expected value) in CCR animals and 88.0 ± 5.8 kJ day^−1^ ind^−1^ (n=8) in controls (117% of the expected value)(F_1,11_=8.92, P=0.012, ANCOVA)(Fig 4a). There were no significant differences in basal metabolic rate (BMR) across seasons and treatments (Fig 4b), but the factorial scope for DEE (DEE/BMR), a measure of the aerobic work capacity, resulted significantly different across seasons and treatments, where in winter control animals had 62% higher value compared with CCR animals (6.45 ± 0.58 over 4.04 ± 0.45, Fig 4c; F_1,11_=5.37, P=0.040, ANOVA)(Fig 4c). Body mass was significantly reduced in CCR animals by 70% during winter compared with their summer values, whereas control individuals did not show seasonal differences (Fig 4d). Summer (pooled: control and CCRs) DEEs were significantly correlated with body mass (R^2^=0.61, P=0.039, n=24, Fig 4d-e), which was maintained in winter, with a difference in intercepts between control and CCR animals (Fig 4f, F_1,13_=8.32, P=0.013, ANCOVA).

**Fig 4.**
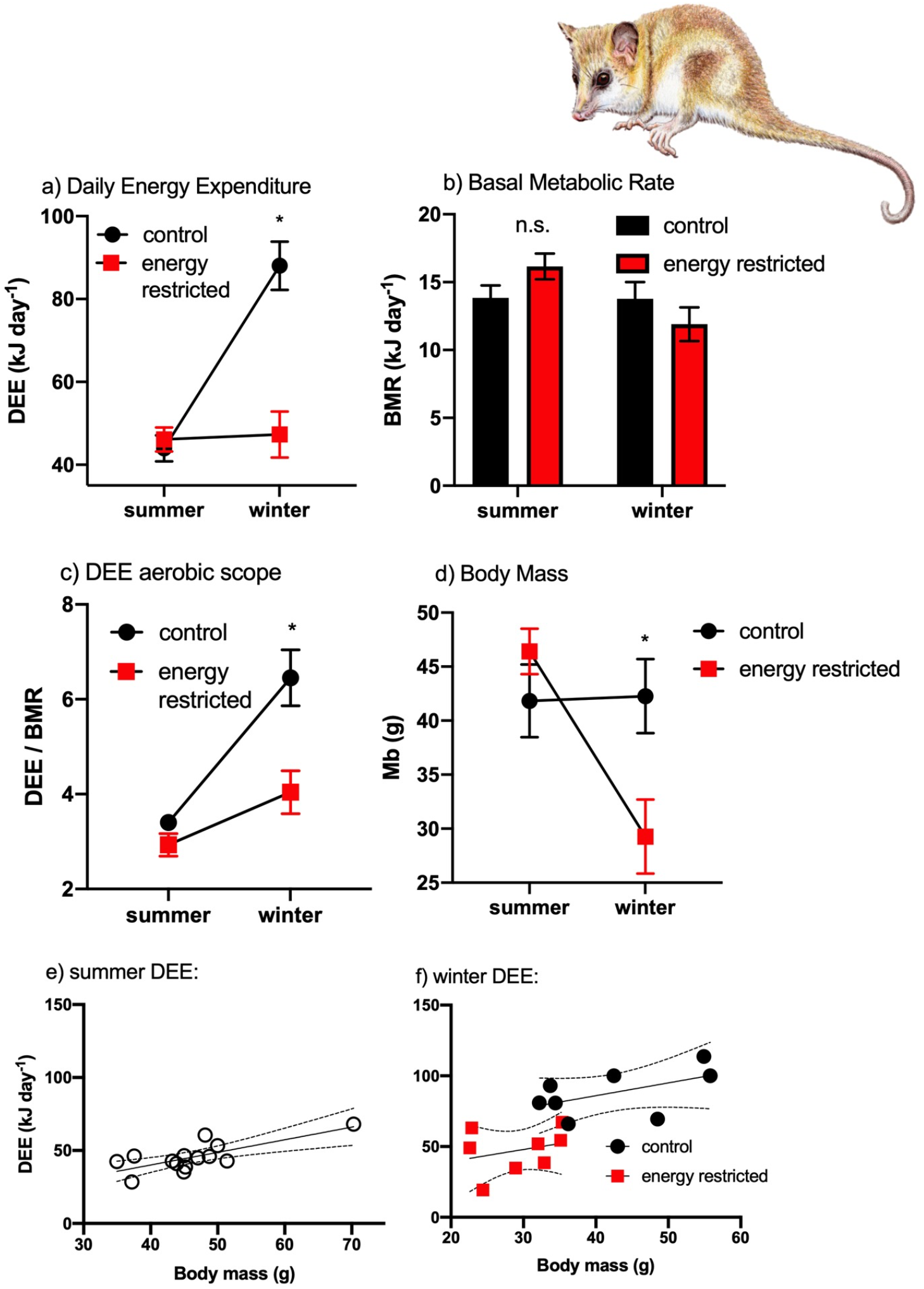
a) Daily energy expenditure (DEE) in summer and winter *D. gliroides* under the CCR and control conditions; b) basal metabolic rate; c) DEE aerobic scope; d) body masses, e) scaling of summer animals for both CCR and control groups pooled; d) scaling of winter animals. Significance (P<0.05) is denoted after a repeated measures ANOVA.

## Discussion

Several authors have calculated the amount of energy saved by specific sections of the hibernation cycle, frequently in a single torpor-arousal cycle and sometimes during multiple events (Geiser 1988, Holloway and Geiser 1995, Schmid and Speakman 2000, Bozinovic et al. 2007, Nespolo et al. 2010, Geiser 2013). These values vary from 99% in single torpor bouts compared with normothermic values, to 15% for multi-day torpor bouts in some hibernators, including the costs of arousals (Wang 1978, Geiser 2004, 2013). However, establishing the precise impact of hibernation on the energy budget of free ranging animals is especially difficult, since a control condition (i.e., a situation without hibernation, keeping all else equal) is hard to obtain. To the best of our knowledge, this has been calculated indirectly on laboratory animals, once in eutherians, the Richarson’s ground squirrel (*Urocitellus richardsonii*, (Wang 1978) and once in a marsupial, in pygmy-possum (*Cercartetus nannus*; (Geiser 2007). Both estimations indicate enormous energy savings by hibernation: 87.7% and 97.5%, respectively, after comparing hibernation energy expenditure with the predicted metabolism of active animals. Our results of daily energy expenditure (DEE) in energy restricted animals and controls provide a direct estimation of this value, with the caveat that during the coldest months (July-October) on average only 69% of CCR animals were in torpor and 25% of controls were in a similar condition. However, these values coincide well with the doubly labelled water method (DLW) estimations, for which all CCR animals were torpid at the moment of sampling, and all control animals were active at this moment. Recalling from Results, *DEE_CONTROL-FOOD_* = 98.6 kJ day^−1^ind^−1^ and *DEE_CONTROL-DLW_* = 88.0 kJ day^−1^ ind^−1^, and averaging, gives *DEE_CONTROL_* = 93.3 kJ day^−1^ ind^−^1. On the other hand, *DEE_CCR-FOOD_* = 48.7 kJ day^−1^ind^−1^ and *DEE_CCR-DLW_* = 47.3 kJ day^−1^ ind^−^1, gives an average *DEE_HIBERNATION_* = 48.0 kJ day^−1^ ind^−1^. This reveals a net hibernation savings of 51.4% (=*DEE_HIBERNATION_ / DEE_CONTROL_*). This smaller value, compared with Belding’s ground squirrel and pigmy possums can be explained by the fact that our *Dromiciops* were experiencing outdoor/field conditions, which includes the thermal impact of natural thermal variations and spontaneous activity bursts during interbout arousals.

According to Humphries et al. (2002)(Humphries et al. 2002) (see also: (French 1985), fat reserves predict wintering hibernation survival, because when “the size of the reserve is less than the rate of depletion times the length of the winter, the hibernator will not survive”. This assertion is true assuming that animals don’t ingest food during hibernation (but see Fig 3). Without eating, a hibernating *D. gliroides* spending 48 kJ day^−1^ ind^−1^ will need 4,320 grams of fat to survive a winter of 90 days (energy content of fat: 39.7 kJg^−1^)(Walsberg and Wolf 1995), which is unrealistic for a 40g animal. It is clear then, that animals regulate food ingestion during interbout arousals, in some way “calculating” torpor incidence for energy management.

Basal metabolic rate (BMR), which is one of the most measured variable in physiological ecology, representing maintenance costs in endotherms (Konarzewski and Diamond 1995, Ricklefs et al. 1996, White and Seymour 2003, McKechnie et al. 2006, Clarke et al. 2010), surprisingly did not vary between seasons or treatments. Instead, the scope for aerobic activity (DEE/BMR), a measure of how hard animals are working when active, showed a significant 89% increase from summer to winter in control animals, but a modest 37.9% increase in CCR animals (from Fig 4d). Thus, CCR animals, in addition of saving energy by hibernation maintained a lower aerobic capacity probably by reducing the amount of metabolically active tissues (Bozinovic et al. 1990, Campbell and MacArthur 1998, Nespolo et al. 2002).

Mueller and Diamond (2001)(Mueller and Diamond 2001) postulated food availability (or net primary productivity) as a unifying factor for explaining adaptive variation in energy expenditure across species, ecosystems, latitude, temperature or rainfall. This idea is related to the more general “pace-of-life” theory of metabolism and life histories, which proposes that populations evolving for a long time at low productivity also evolve low levels of energy expenditure (Wikelski et al. 2003, Careau et al. 2010, Le Galliard et al. 2013, Londono et al. 2015, Pettersen et al. 2016). Our results support the idea that hibernation represents a “pace-of-life” adaptation to environments characterized by seasonal reductions of primary productivity (i.e., characteristics of temperate regions), where hibernation acts a physiologically regulated metabolic switch-off coupled with the period of low primary productivity (winter){Turbill, 2011 #3341}. In this sense, the fact that hibernation is present in several unrelated species living in the same environments supports the view of hibernation as a convergent feature of mammals (Boyles et al. 2013). In fact, *D. gliroides*, the only South American (SA) mammal described as a hibernator (Bozinovic et al. 2004), has a distribution range in South America between 35° and 45° S, a narrow latitudinal strip that in the Southern hemisphere includes a few landmasses (the tip of South Africa, Southern Australia including Tasmania and most part of New Zealand). This contrasts with the vast extensions of territories included in this range at the Northern hemisphere, from which almost all hibernating species have been identified (Humphries et al. 2002, Boyles et al. 2008, Ruf and Geiser 2015). Perhaps the right terrestrial environment at the Southern hemisphere simply did not provide enough land area for hibernation to evolve more frequently.

Mesocosm studies (i.e., outdoor experiments examining natural environments under controlled conditions) provide a fundamental link between field surveys and laboratory experiments (Kennedy 1995, Verdier et al. 2014, Kurz et al. 2017, Maugendre et al. 2017, Scharfenberger et al. 2019). However, they are particularly scarce in ecological physiology (however, see references (Merritt et al. 2001, Levy et al. 2012, Gao et al. 2015), a field with a long tradition on laboratory work (see ref (Humphries et al. 2003b) and cited references). We encourage more of such experiments. Researchers will surprise how simple and cost-effective they are, as one single long-term experiment could replace many small laboratory trials.

## Acknowledgements

This work was funded by FONDECYT grant 1180917 to RFN, ANID PIA/BASAL FB0002 to FB, RFN and PS. We also thank Enrico Rezende for a critical review of the manuscript.

## Supplementary Material and Methods

### Enclosures

Each enclosure had a internal volume of 2 m^3^, and was manufactured in zinc by a large 1.8-diameter cylinder buried 10 cm in the ground, which gave a 0.8 m height above ground. Each ceiling was framed in timber, and had a mesh that allowed the entrance of light and humidity, but avoided the escape of the animals or predator’s attack. Then we included a tri-dimensional arrangement of *Nothofagus* twigs and logs, native bamboo (*Chusquea quila*) in each enclosure, and the floor was covered by mosses and bamboo leaves, which are known to be essential for *D. gliroides* nests building (Hershkovitz 1999, Honorato et al. 2016), resembling forest conditions (see Fig 1b-d in main text). We also included one removable hibernaculum per enclosure, which consisted in a hollowed log of about 30×10×15 cm, cut longitudinally that was put over the ground in a way that allowed animals to enter, cluster, rest, or hibernate. Each hibernaculum was sealed at each end by a timber cover with a small hole in the middle, to allow animal entrance. In each enclosure, we also put one max/min thermometer, one temperature data logger (HOBO®) for continous T°C recording and water ad libitum.

### Diet preparations

*D. gliroides* is an omnivorous marsupial with well-known dietary preferences (Cortes et al. 2011, Rodriguez-Cabal and Branch 2011, Contreras et al. 2014), thus we offered three dietary items to them in separate plates: apple compote, canned tuna (in water) and blend (i.e., equal parts mix between berry jam and baby cereals plus 50% of water) (Contreras et al. 2014))(see Fig 1e in main text). We also added a polyvitamin mixture in the diets (0.3 mg kg^−1^ inveade®). The apple compote and the tuna were offered as they are obtained from the commercial suppliers. We always used the same commercial suppliers. Three samples of each diet were dried and calorimetrically analyzed in a Parr calorimeter (Illinois, USA), showing similar energy contents (dry weight)(tuna: 23.04 ± 3.4 kJg^−1^; blend: 17.90 ± 0.12 kJg^−1^; apple compote: 15.89 ± 0.48 kJg^−1^)(see details in Table S1). We calculated food consumption using marsupial allometric equations (Nagy 2001) and considering a maximum energy expenditure that is six times basal metabolic rate (Bozinovic et al. 2004, Nespolo et al. 2010, Franco et al. 2012).

**Table S1.** Nutrient content of the experimental diets provided to the enclosures. Each enclosure received three dietary items: (1) a homogenized blend of jam and cereal diluted in 50% water, (2) a weighed amount of tuna and (3) a weighed amount of apple compote from a commercial supply (see methods for details).

| **Commercial label** | **Jam** | **Cereal** | **Canned**<br>**tuna** | **Apple**<br>**compote** |
| --- | --- | --- | --- | --- |
| Calories (KJ/100g) | 887 | 1,564.8 | 280.3 | 281.2 |
| Protein (%) | 0.3 | 9 | 15 | 0.3 |
| Total fat (%) | 0.2 | 1.8 | 0.4 | 0.3 |
| Total Carbohydrate (%) | 52.2 | 80.5 | 0.5 | 16 |
| Total sugars (%) | 51.7 | 26.0 | 0.5 | 16 |
| Sodium (mg/100g) | 13 | 80 | 314 | 4 |

### Thermographic imaging

For characterizing thermoregulatory abilities of hibernating *D. gliroides*, we visited the enclosures every week, uncovered each hibernaculum, took a digital photo and an infra-red photograph of clustered torpid individuals using a thermograph (FLIR systems, Oregon, USA) set for an emissivity of 0.98 (Fig. 1f-g, total images: 328). This infrared imaging permitted us to measure in situ external body temperatures (*T_TORPID_*), by averaging the temperature of a polygon drawn of the image of each animal using the FLIR tools software. We also measured the mean temperature of the substrate 10 cm apart of the cluster (*T_SUBSTR_*). With this information, we calculated the thermal differential (*T_DIFF_* = *T_TORPID_* – *T_SUBSTR_*) for each animal, which is a measure of heat conservation in torpor. After recording these images, we measured cloacal temperature on each animal, using a Cole-Parmer copper-constantan thermocouple inserted 1 cm in the cloaca. This record was obtained within a few minutes after taking the images (otherwise it was discarded). Cloacal temperature was correlated with *T_TORPID_* (R^2^ = 0.68; P < 0.01; Fig. S1, n= 410). Finally, each torpid animal was weighed and released back in the hibernaculum. We also recorded the size of the cluster and whether they were found within the hibernaculum. We also classified each animal as torpid or active by visual inspection (see Fig 1h in main text).

**Fig S1.**
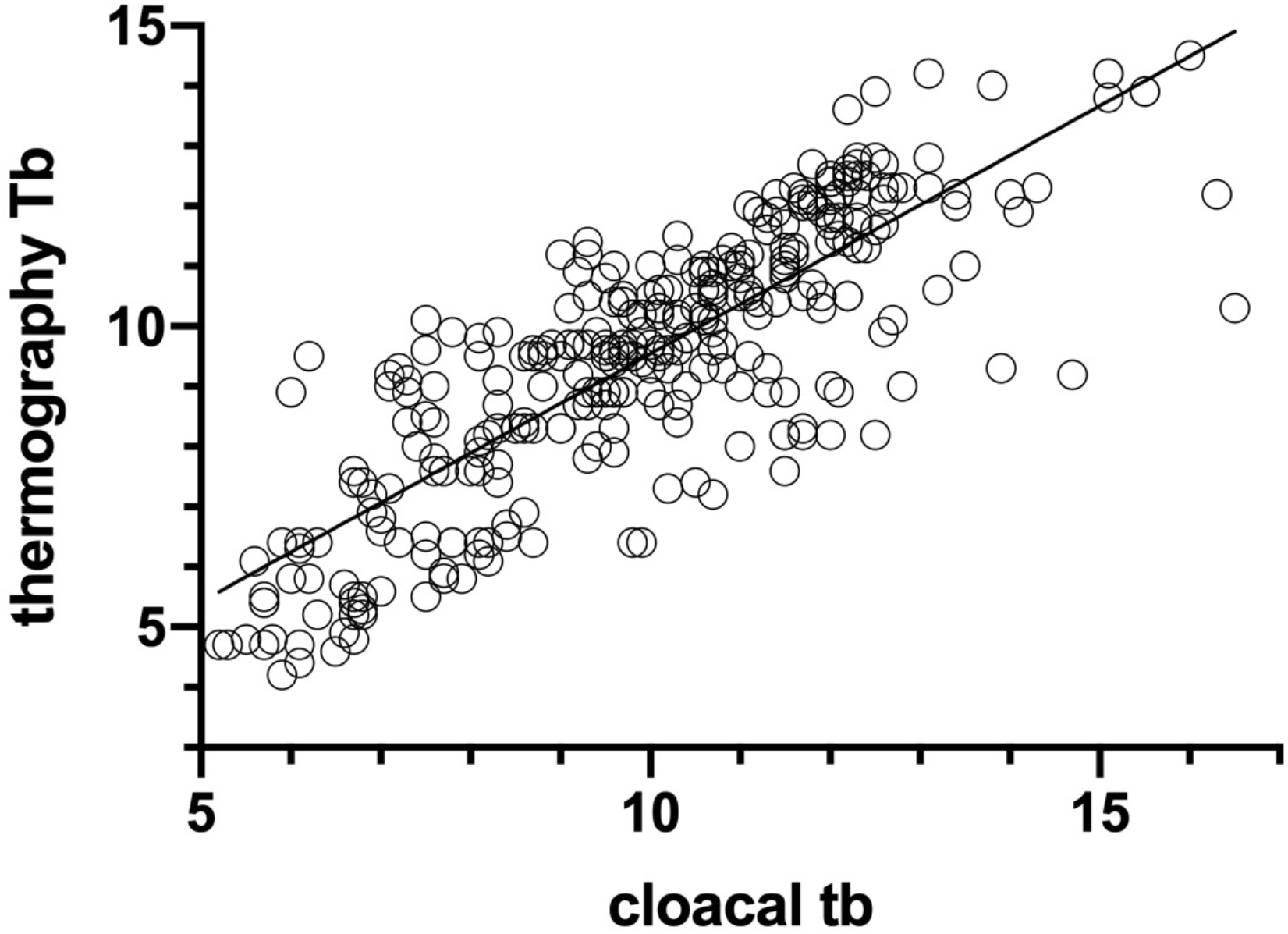
Bivariate relationship between surface skin temperature measured by thermographic images and cloacal temperature, measured by a copper-constantant thermocouple, in each animal.

### Doubly labelled water

This method has been previously validated by comparison to indirect calorimetry in a range of small mammals (e.g. Speakman and Krol, 2005). A weighed amount of DLW was injected intraperitoneally into each individual. A blood sample (100ul) was collected from the tail vein into glass capillaries and flame sealed 1 and 48 hours later. Background samples were collected from some individuals prior to dosing. Analysis of the isotopic enrichment of blood was performed blind using a Liquid Isotope Water Analyser (Los Gatos Research, USA) (Berman et al. 2012). Initially the blood encapsulated in the capillaries was vacuum distilled (Nagy 1983), and the resulting distillate was used. Samples were run alongside three lab standards for each isotope and International standards to correct delta values to ppm. Equation 7·17 of Speakman (1997)(Speakman 1997) assuming a single-pool model was used to calculate rates of CO_2_ production as recommended for use in animals less than 1 kg in body mass (Speakman 1997). There are several approaches for the treatment of evaporative water loss in the calculation (Visser and Schekkerman 1999). We assumed evaporation of 25% of the water flux (equation 7.17: Speakman 1997) which minimizes error in a range of conditions (Visser and Schekkerman 1999, Van Trigt et al. 2002). CO_2_ production was converted to DEE using the Weir equation (Weir 1990).

### Basal metabolic rate

Briefly, metabolic rate was recorded using a LiCor 6251 CO_2_ analyzer in a 1L metabolic chamber and a flow rate of 1,000 ml min^−1^, after scrubbing water and CO_2_ from the incoming air. The metabolic chamber was located in an incubator, and ambient temperature was set to thermoneutrality (30°C) which was continuously recorded by a thermocouple located inside the incubator. These measurements were completed after a day of acclimation to the laboratory and after food had been removed for 8 hrs. Metabolic trials all took place during the typical rest phase of the animals (between 8am and 7pm). Each measurement had a duration of three hours and most animals slept after the first hour in the chamber, which was checked by visual inspection though a small window in the incubator. BMR (mlCO_2_ h^−1^) was calculated from the three lowest steady-state values during the last 30 min of recording, and converted to kJ assuming an RQ=0.71 (Walsberg and Wolf 1995).

### Statistical analyses

We fitted Mixed-Effects Generalized Linear Models (GLMM) with a gaussian error distribution and an ‘identity’ link function on the previously defined variables. We included individual ID, enclosure, and sampling week as random effects to account for inter-individual and inter-enclosure variability, along with the repeated measures in time (Zuur et al. 2009). To estimate the best explanatory variables for torpor occurrence, we fitted a GLMM with a binomial error distribution and a ‘logit’ link function (Beckerman et al. 2017), including treatment, body mass, and group size as predictors (fixed effects) and individual ID, enclosure and sampling week as random effects, as described above. To explore the factors that influence heat conservation in torpid animals, we fitted additional models using the same parameters on a subset of data of torpid animals. Then, we fitted one more GLMM to assess the factors determining *T_DIFF_*, using CCR treatment, body mass, and group size as predictors (fixed effects) and individual ID, enclosure and sampling week as random effects, as previously described. We estimated GLMM parameters and their significance using a restricted maximum likelihood approach with a Kenward-Roger approximation to estimate degrees of freedom (Halekoh and Hojsgaard 2014). We performed all analyses using R 3.6.0 (Team 2019), with the packages mgcv (Wood 2011), lme4 (Bates et al. 2013), lmerTest (Kusnetzova et al. 2015), pbkrtest (Halekoh and Hojsgaard 2014), and ggplot2 (Wickham 2016).

## Supplementary Results

### Thermoregulation during torpor

As soon as ambient temperature fell below ~12°C, we observed packed clusters of torpid animals, sometimes within a compact nest of interwoven leaves of native bamboo (*Chusquea quila*) and mosses, or sometimes just buried in the ground. However, thermoregulatory adjustments during hibernation between CCR and control animals were not different, as revealed by thermographic images (summarized in Fig. S2 and Table S2, n= 328), and by the frequency of clustering or hibernacula use (summarized in Fig. S3, n=530 and 618, respectively). Although the GLMM model using torpor occurrence as a binomial variable showed several significant effects of the CCR treatment, indicating complex interaction among food deprivation, cluster size and body mass (Table S2), there were non-significant effects of these variables on the thermal differential between animals and substrate, estimated by the analysis of *T_DIFF_* (Table S3). Thermoregulatory variables such as the T_B_/T_A_ slope comparison between control and CCR (Fig. S2a-c) and the comparison of slopes of the logistic regression of torpid and active animals (Fig. S2d; n=795 and 342, control and CCR respectively) were non-significant. Also, the most frequent substrate temperature for torpor in control individuals (median=10.05, min=4.8, max=16.2°C, n=130) was nearly identical with CCR individuals (median=10.1, min=4.2, max=15.9°C, n=148, Fig. S2e-f, non-significant differences after a median test). Behavioral strategies for heat conservation such as clustering (control animals formed small groups during torpor, whereas CCR animals did not show any trend, Fig. S3a-b), and hibernacula use (control animals were preferably found within hibernacula, both active and torpid, Fig. S3c-d) indicated absence of behavioral strategies for heat conservation in CCR. In other words, ad libitum fed animals preferred hibernacula irrespectively of being active or torpid.

**Fig S2.**
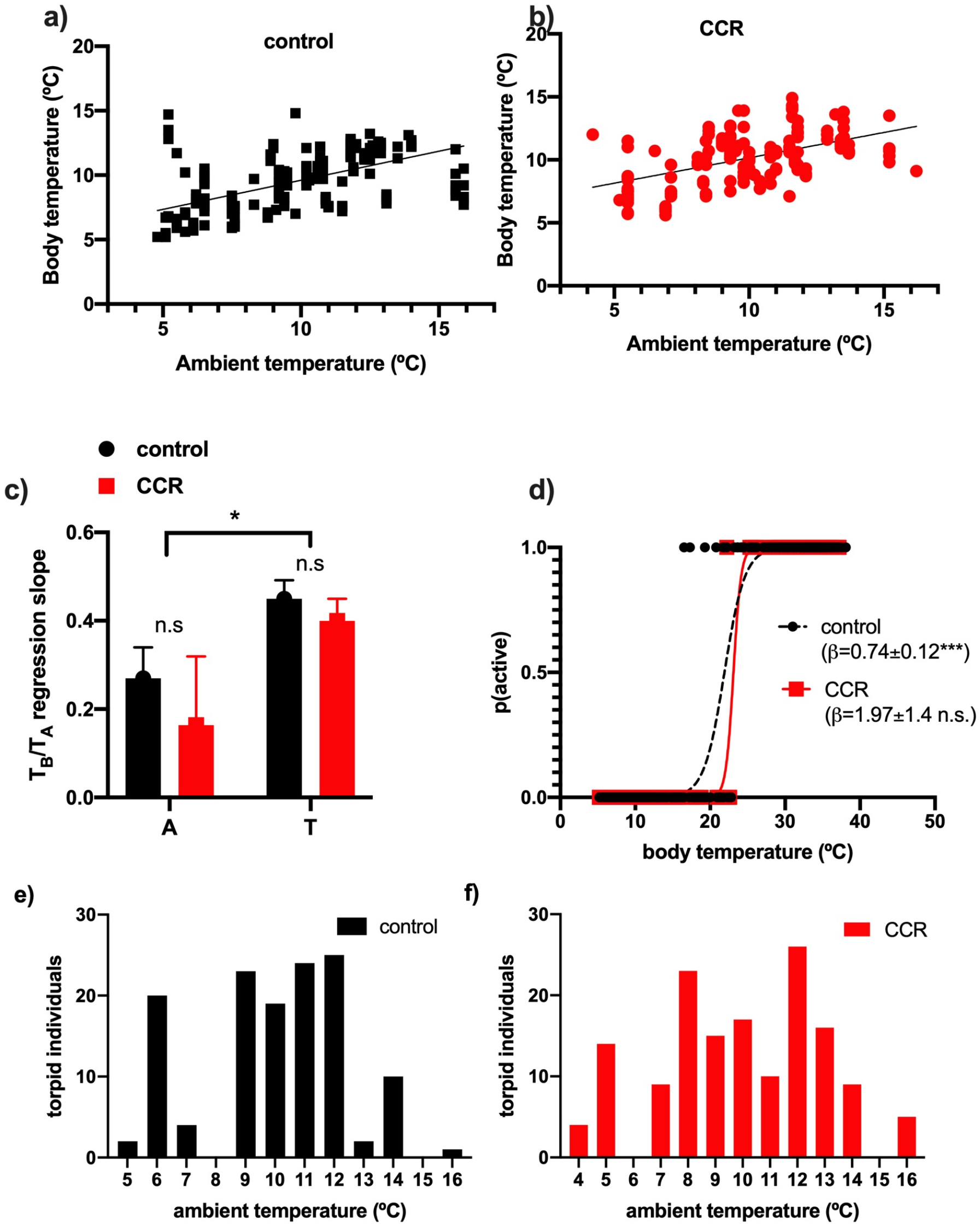
Thermal physiology of D. gliroides under control and energy restricted conditions. Linear regressions (a: control; b: treatment) between ambient temperature and body temperature measured weekly as cloacal temperature in a semi-natural experiment of chronic caloric restriction. Figure S2c shows a comparison of the T_A_/T_B_ slopes (**A**: active; **B**: torpid) calculated above, showing significant differences only for torpid and active individuals: comparisons either within control (F_1,263_=19.9; P=0.018; ANCOVA homogeneity of slopes model) or within energy restricted animals (F_1,387_=19.7; P=0.018; ANCOVA homogeneity of slopes model). Non-significant differences were found for control/treatment comparisons within torpid or within active animals (indicated). Figure S2d) logistic regression between body temperature and probability of being active, showing a rewarming threshold in T_B_ of about 22°C, but it was non-significant for energy restricted animals. Figure S2e and f) shows substrate preferred temperatures in control and energy restricted individuals. Both distributions have identical medians (=10.1°C).

**Fig S3.**
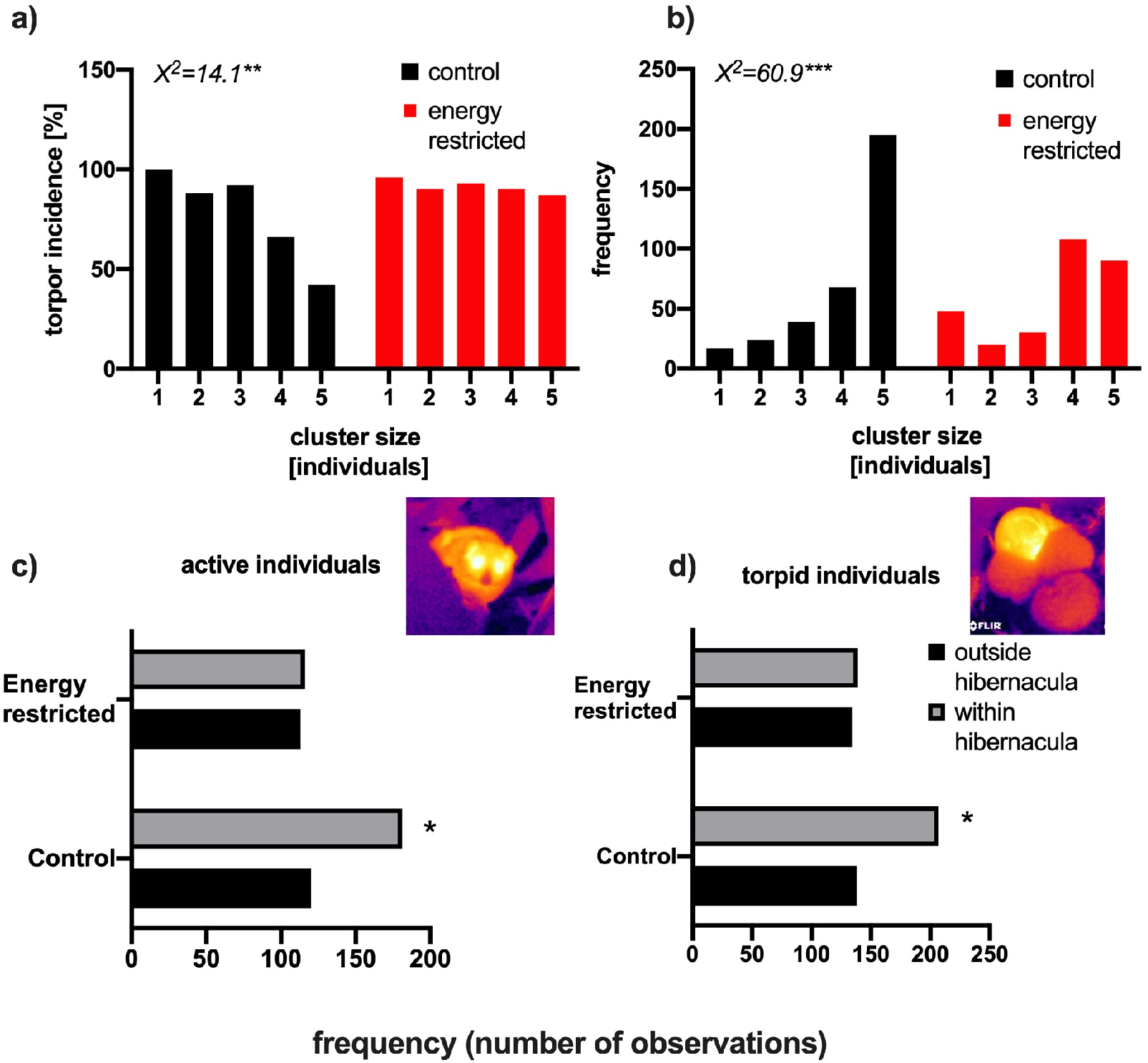
Frequency distributions of animals forming groups or using hibernacula during the CCR experiment. a) torpor incidence in function of cluster size; b) total frequency of cluster size; c) hibernacula use in active animals and d) hibernacula use in torpid animals. Significant values indicating different frequencies across categories, are indicated after a chi-square contingency table (indicated in the figure) and Fisher exact test (*P<0.001).

**Table S2.** Results of a generalized linear mixed model fit by restricted maximum likelihood, for the binomial response variable “status” (active/torpid) using the logit link (n= 795). The model was: status ~ treatment (restricted/control) + body mass (M_B_) + cluster size (1-5 individuals) + enclosure (random factor) + ID (random factor) + week (random factor). The model included all possible interactions.

| Variable | Estimate | SE | z-value | P-value |
| --- | --- | --- | --- | --- |
| (Intercept) | 10.220 | 4.169 | 2.451 | 0.014 |
| caloric restriction treatment | -21.469 | 6.640 | -3.234 | 0.001 |
| mass | -0.485 | 0.134 | -3.621 | <0.001 |
| group.size | -1.889 | 0.842 | -2.243 | 0.025 |
| dietTreatment:mass | 0.652 | 0.202 | 3.225 | 0.001 |
| dietTreatment:group.size | 3.092 | 1.468 | 2.106 | 0.035 |
| mass:group.size | 0.091 | 0.027 | 3.301 | 0.001 |
| dietTreatment:mass:group.size | -0.114 | 0.046 | -2.496 | 0.013 |

**Table S3.**
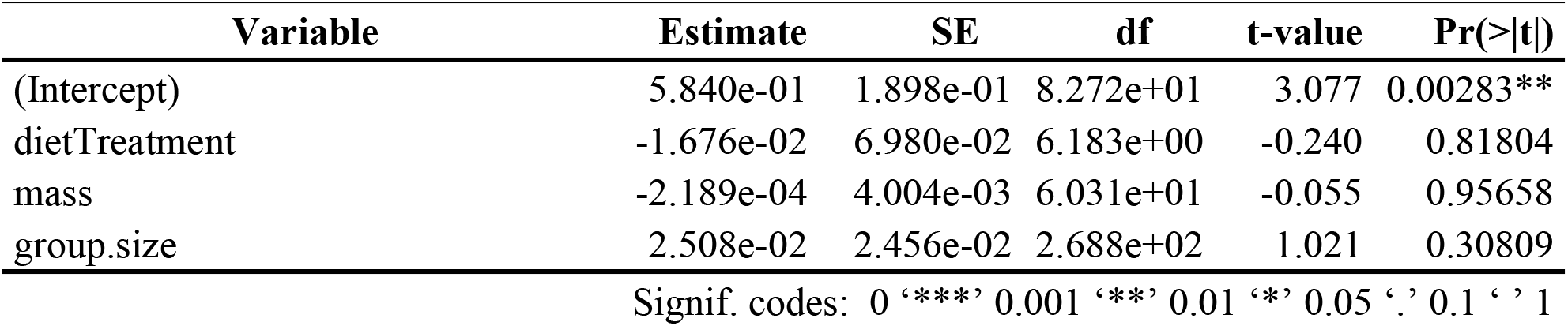
Results of a generalized linear mixed model fit by restricted maximum likelihood, for the response variable “T_DIFF_” (thermal differential), obtained using thermographic pictures in clustered hibernating animals (n= 328). The model was: T_DIFF_ ~ treatment (restricted/control) + body mass (M_B_) + cluster size (1-5 individuals) + enclosure (random factor) + ID (random factor) + week (random factor). The model included all possible interactions.

| Variable | Estimate | SE | df | t-value | Pr(>\|t\|) |
| --- | --- | --- | --- | --- | --- |
| (Intercept) | 5.840e-01 | 1.898e-01 | 8.272e+01 | 3.077 | 0.00283** |
| dietTreatment | -1.676e-02 | 6.980e-02 | 6.183e+00 | -0.240 | 0.81804 |
| mass | -2.189e-04 | 4.004e-03 | 6.031e+01 | -0.055 | 0.95658 |
| group.size | 2.508e-02 | 2.456e-02 | 2.688e+02 | 1.021 | 0.30809 |

Signif. codes: 0 ‘\*\*\*’ 0.001 ‘\*\*’ 0.01 ‘\*’ 0.05 ‘.’ 0.1 ‘ ’ 1

## Notes

### Competing Interest Statement

The authors have declared no competing interest.

